# An autoimmune disease risk variant has a *trans* master regulatory effect mediated by IRF1 under immune stimulation

**DOI:** 10.1101/2020.02.21.959734

**Authors:** Margot Brandt, Sarah Kim-Hellmuth, Marcello Ziosi, Alper Gokden, Aaron Wolman, Nora Lam, Yocelyn Recinos, Veit Hornung, Johannes Schumacher, Tuuli Lappalainen

## Abstract

Functional mechanisms remain unknown for most genetic loci associated to complex human traits and diseases. In this study, we first mapped *trans*-eQTLs in a data set of primary monocytes stimulated with LPS, and discovered that a risk variant for autoimmune disease, rs17622517 in an intron of *C5ORF56*, affects the expression of the transcription factor *IRF1* 20 kb away. The cis-regulatory effect on *IRF1* is active under early immune stimulus, with a large number of *trans*-eQTL effects across the genome under late LPS response. Using CRISPRi silencing, we showed that the SNP locus indeed functions as an *IRF1* enhancer with widespread transcriptional effects. Genome editing by CRISPR further indicated that rs17622517 is indeed a causal variant in this locus, and recapitulated the LPS-specific *trans*-eQTL signal. Our results suggest that this common genetic variant affects inter-individual response to immune stimuli via regulation of *IRF1*. For this autoimmune GWAS locus, our work provides evidence of the causal variant, demonstrates a condition-specific enhancer effect, identifies *IRF1* as the likely causal gene in *cis*, and indicates that overactivation of the downstream immune-related pathway may be the cellular mechanism increasing disease risk. This work not only provides rare experimental validation of a master-regulatory *trans*-eQTL, but also demonstrates the power of eQTL mapping to build mechanistic hypotheses amenable for experimental follow-up using the CRISPR toolkit.

## Introduction

The discovery of tens of thousands of genetic loci associated to complex diseases and traits has introduced the challenge of characterizing the biological mechanisms that mediate these associations. Most GWAS loci include a large number of associated variants in linkage disequilibrium (LD) and are located in noncoding regions with likely gene regulatory functions. Therefore, it is typically unknown what is the causal variant, affected regulatory element, target gene in *cis*, and downstream molecular pathways that mediate genetic associations to complex disease phenotypes. Furthermore, the cellular context where these effects take place is often elusive, with increasing evidence that effects can be not only specific to cell types, but also cell states.

A key approach to answer these questions is expression quantitative trait locus (eQTL) analysis to discover associations between genetic variation and gene expression. Most widely applied in *cis*, eQTLs across diverse tissues have been shown to be strongly enriched for GWAS signals and have the ability to pinpoint potential target genes in GWAS loci (Gamazon et al., 2018). Furthermore, response *cis*-eQTL mapping for immune cells with *in vitro* stimuli have provided powerful evidence of genetic regulatory effects that are not only specific to tissues but also to cell states, and disease associations that are driven by disrupted response to environmental stimuli (Fairfax et al., 2014; Kim-Hellmuth et al., 2017; Lee et al., 2014; Nédélec et al., 2016). *Trans*-eQTL mapping for distant, typically interchromosomal, genetic regulatory effects has the additional potential for elucidating regulatory pathways of the cell, and *trans*-eQTLs have been shown to be very strongly enriched for GWAS signals (Aguet et al., 2019; Small et al., 2018; Võsa et al., 2018; Westra et al., 2013). Unfortunately, robust discovery of *trans*-eQTLs has been challenging due to the large multiple testing burden, smaller effect sizes, and higher tissue-specificity as compared to *cis*-eQTLs (Aguet et al., 2019). While a large number of *trans*-eQTLs are mediated by a *cis*-eQTL, indicating potential biological mechanism, *trans*-eQTLs in humans rarely have strong master-regulatory effects on pathways of multiple genes (Aguet et al., 2019; Battle et al., 2014; Grundberg et al., 2012; Westra et al., 2013).

Despite the power of eQTL characterization, the approach has limitations that open up possibilities for complementary experimental analysis. Association studies cannot provide definitive evidence for distinguishing causal regulatory variants from their LD proxies, creating the need for massively parallel screens and CRISPR assays to test whether specific genetic perturbations indeed affect gene regulation. Furthermore, experimental characterization and validation of eQTL associations has been sparse, which is a problem especially for *trans*-eQTLs that do not always replicate well between independent data sets (GTEx Consortium et al., 2017). Finally, with the functional genomics toolkit now including eQTL mapping and diverse experimental approaches in cellular models, understanding the transferability of *ex vivo* eQTL results and cellular models is poorly known. Understanding these aspects is a particularly burning question for experimental follow-up of disease-associated loci.

In this study, we mapped *trans*-eQTLs in a monocyte data set with LPS stimulus time points, with *cis*-eQTL discovery reported in earlier work (Kim-Hellmuth et al., 2017). We discovered an inflammatory bowel disease (IBD) GWAS variant rs17622517 that affects the expression of the transcription factor *IRF1* in *cis* under early immune stimulus, and a large number of genes in *trans*-under late stimulus. Using CRISPRi silencing and CRISPR genome editing, we show that the SNP locus indeed functions as an enhancer, identify the causal variant of this association, and demonstrate that the LPS-specific *trans*-eQTL signal can be recapitulated in a cellular model. Our results suggest that this common genetic variant affects inter-individual response to immune stimuli via regulation of *IRF1*, providing a strong hypothesis for the functional mechanism of the IBD risk association in this locus.

## Results

### eQTL discovery and characterization in cis and trans

We mapped response eQTLs in *cis* in primary monocytes under baseline and under the innate immune stimulus with LPS, using a data set of 134 donors with SNP genotyping and gene expression measured under baseline, and early (90 mins) and late (6h) LPS treatment that triggers the Toll-like receptor 4 (TLR4) pathway. The *cis-*eQTL discovery is described in (Kim et al., 2014; Kim-Hellmuth et al., 2017), and here our focus was how early immune response *cis*-eQTLs may translate to later *trans*-eQTL effects. To this end, for the lead variants of the 126 *cis*-eQTLs at 90 min, we mapped *trans*-eQTLs for all the expressed probes at 6h. We discovered 47 probes with at least one *trans*-eQTL (*trans*-eGenes) at 5% false discovery rate (FDR) and 204 probes at 25% FDR, with 69 *cis*-eQTLs having at least one *trans* effect at 25% FDR (Supplementary Table 1).

One locus stood out by being associated to a large number of *trans*-eGenes: 12 at 5% FDR and 232 at 50% FDR - substantially more than the other loci (Fig. 1a). Interestingly, the *trans*-eQTL is a cis-eQTL for the nearby *IRF1* only under early LPS response (Fig. 1b), and the *trans-* eGenes are enriched for *IRF1* target genes, defined as having an *IRF1* motif within 4 kb of their transcription start site (TSS) according to MSigDB (p = 1.8e-4). These *trans-*eQTLs are mostly active only at 6 hours after LPS stimulation (Fig. 1c). This pattern suggests a hypothesis where early LPS stimulation activates the *cis-*eQTL effect on *IRF1* first, which then affects the expression of downstream direct and indirect targets of *IRF1*. The alternative allele of the lead variant rs17622517 is associated with upregulation of *IRF1* upon early immune stimulus, and upregulation of 11 out of 12 of its strongest trans-eGenes as well (Supplementary Figure 1).

**Figure 1.**
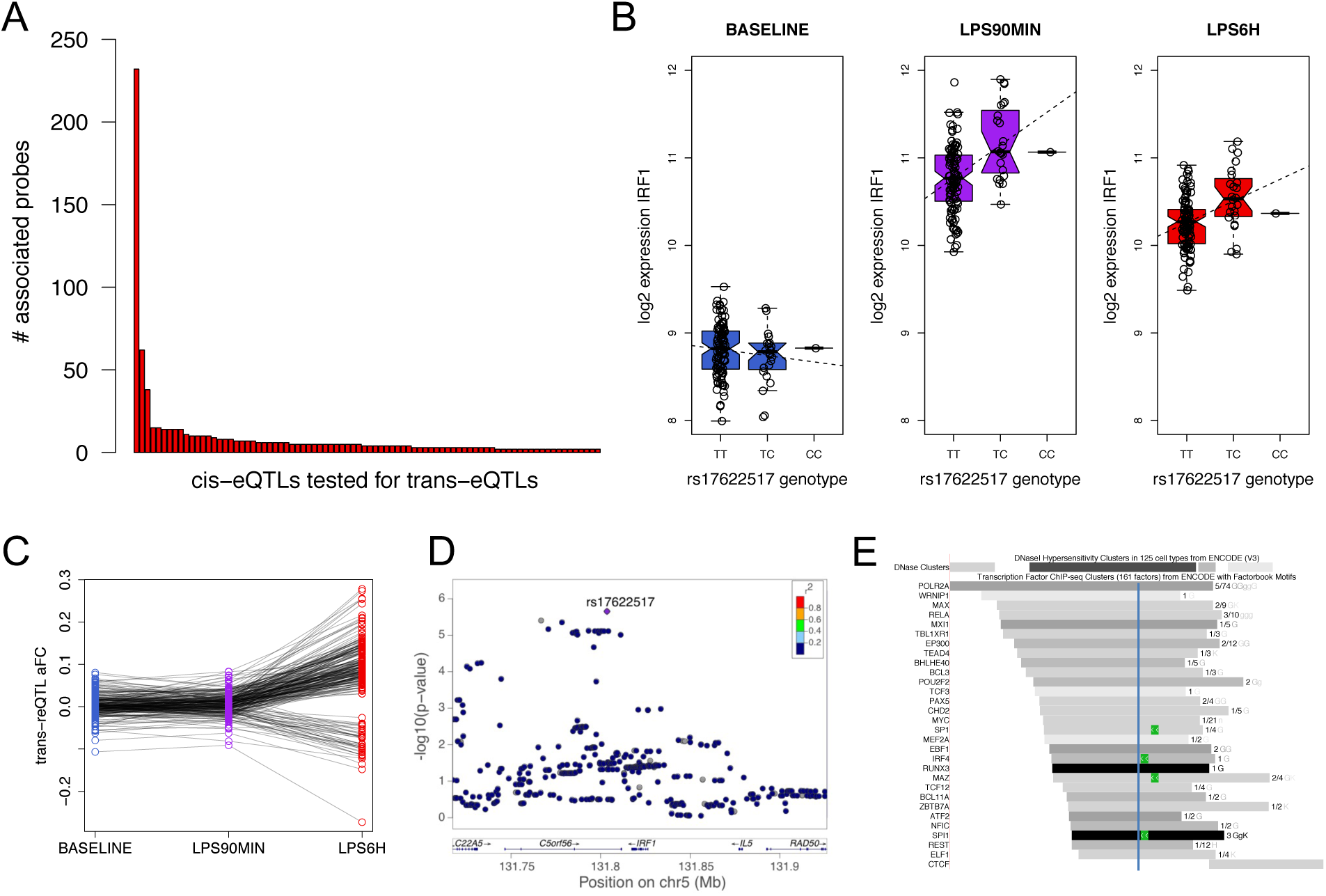
eQTL analysis. a) The number of trans-eGenes at 0.5 FDR per each tested locus with 6h LPS stimulation. The leftmost bar is the IRF1 locus; b) the rs17622517 cis-eQTL for IRF1, and c) and its trans-eQTL effect sizes (allelic fold change) under the three conditions; d) the association landscape for the IRF1 cis-eQTL signal; e) overlap of rs17622517 (blue line) with transcription factor ChIP-seq peaks (grey) and motifs (green).

Next, we sought to further characterize the eQTL, its likely causal variant and its *cis*-regulatory mechanism. The lead variant rs17622517 is located 23 kb downstream of the *IRF1* TSS in an intron of the gene *C5ORF56* (Fig. 1d). It has no LD-tagging variants, and the other neighboring variants that are also significantly associated with *IRF1* in *cis* (Supplementary Table 2) represent an independent eQTL for *IRF1*, with a lead variant rs147386065 that has a uniform effect under baseline and the two stimulus timepoints (Supplementary Figure 2a). The trans-eQTL signal is significant only for rs17622517, but its effect sizes for the 25% FDR trans-eGenes are correlated to much weaker effect sizes from rs147386065 (Supplementary Figure 2b), suggesting that both genetic effects on *IRF1* may contribute to downstream regulatory changes. For the lead eQTL, rs17622517 is a good candidate for the causal variant since it has no LD proxies, and its genomic context lends further support to this: In ENCODE data, the SNP locus overlaps open chromatin in many cell types, including monocytes (Thurman et al., 2012), and a H3K27Ac peak in the GM12878 lymphoblastoid cell line suggests an active immune cell enhancer in this region. It also overlaps binding sites of numerous transcription factors in lymphoblastoid cell lines and is close to the motifs of many of them, including transcription factors activated in immune response, like IRF4 and RELA, a subunit of NF-κB (Fig. 1e).

Interestingly, this locus has a robust multi-trait genome-wide association signal for inflammatory bowel diseases (IBD; Crohn’s disease, and ulcerative colitis) and ankylosing spondylitis (de Lange et al., 2017; Ellinghaus et al., 2016; Huang et al., 2017). At least three independent causal GWAS signals appear to exist in the locus, of which two match the two independent eQTLs in this locus (de Lange et al., 2017; Ellinghaus et al., 2016). The causal gene(s) in this locus have been unclear, with previous eQTL and other evidence pointing to several potential causal genes (de Lange et al., 2017; Ellinghaus et al., 2016; Huang et al., 2017). While *IRF1* is a plausible candidate as a key immune regulator with links to many ulcerative colitis-related genes (Ciorba et al., 2010; Shi et al., 2019; Wu et al., 2014), functional data implicating *IRF1* has been thus far lacking.

Our findings suggest that this GWAS association for IBD may be driven by a genetic effect that affects response to LPS via a cis-regulatory effect on *IRF1* followed by trans-regulatory effect on its downstream pathway. With a good candidate for a causal variant in a putative enhancer, we next sought to characterize and validate these effects experimentally using enhancer and promoter silencing by CRISPRi and genome editing by CRISPR.

### CRISPRi characterization of the enhancer function and downstream immune response

First, we had to select a cellular model. With monocyte-like THP1 cells being difficult to grow and edit, we chose to use a modified HEK293 cell line HEK293/hTLR4A-MD2-CD14 (Invivogen) co-transfected with the human TLR4A, MD2 and CD14 genes, conferring NF-kB nuclear translocation in presence of Gram-negative lipopolysaccharide (LPS) (Jiang et al., 2005; Poltorak et al., 1998; Shimazu et al., 1999). By RNA-sequencing after LPS stimulus under 4 concentrations and 7 timepoints we verified that the cell line responds to LPS stimulus and activates relevant pathways (Supplementary Figure 3). The transcriptional response appeared slightly delayed compared to primary monocytes, and thus we used 90 minutes and 12 hours post-LPS for early and late immune stimulus time points, respectively.

Next, we studied whether the rs17622517 locus indeed appears to be an enhancer for *IRF1*. To this end, we used CRISPRi with transfection of the dCas9-KRAB-MeCP2 construct (Yeo et al., 2018) with a gRNA targeted to the variant locus to silence its activity (Fig 2a). We also used the same construct to silence the *IRF1* promoter with gRNA targeting 14 bp downstream of the *IRF1* TSS, in order to establish a positive control for *IRF1* silencing. As a CRISPRi negative control we used a gRNA targeting GFP. After transfection with CRISPRi constructs, we treated four replicates of each transfection with LPS for 0h, 90m or 12h, performed RNA-sequencing on the 36 samples with a median of 17 million reads, and analyzed gene expression patterns.

**Figure 2.**
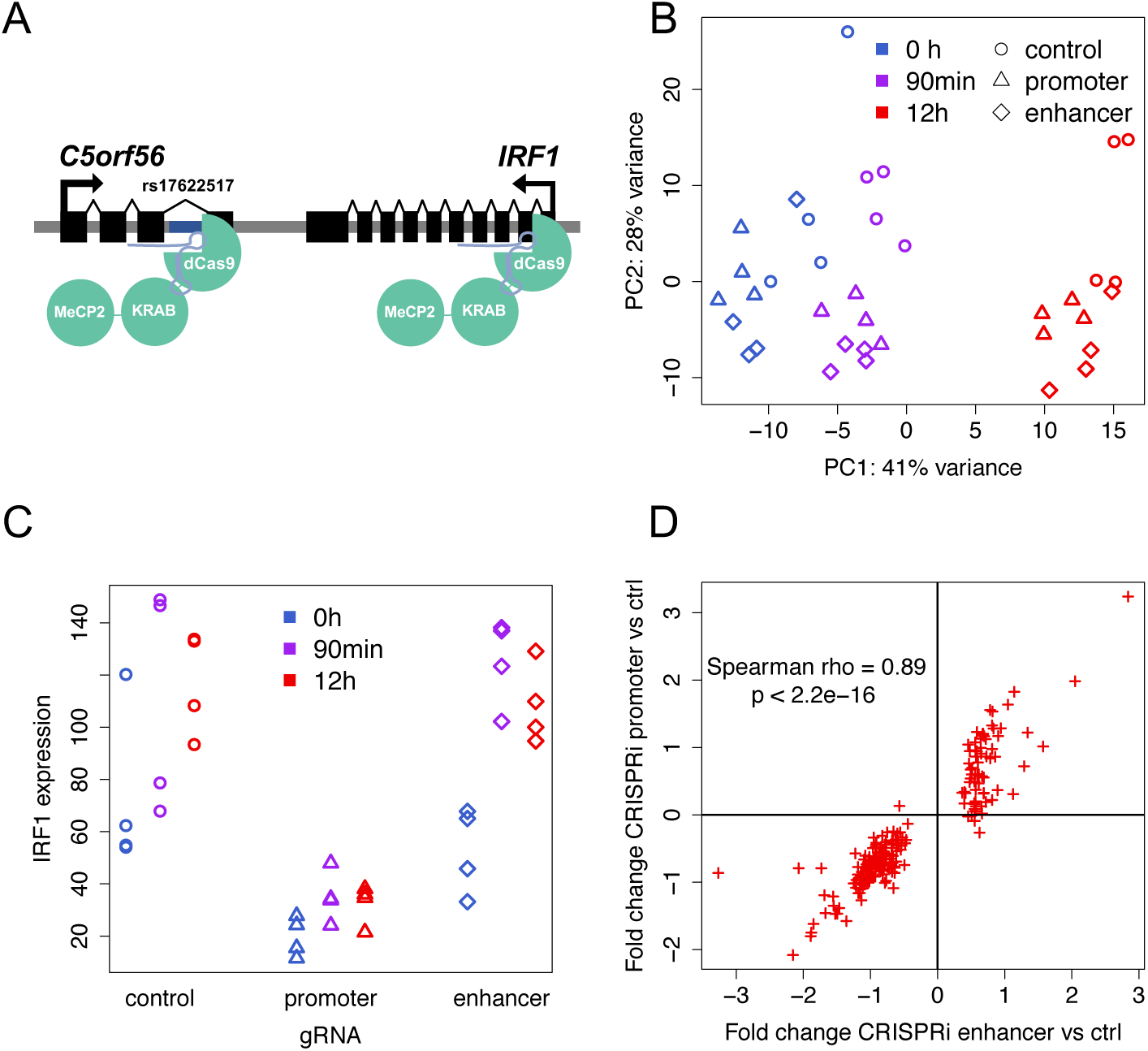
CRISPRi silencing. a) Illustration of CRISPRi silencing of the IRF1 promoter (right) and the putative enhancer locus at rs17622517 (left); b) principal component analysis of gene expression under the different LPS and CRISPRi conditions; c) IRF1 expression (gene counts normalized to sample read depth) under the different conditions; d) The correlation of expression fold changes for promoter and enhancer silencing at 12h LPS, shown for 225 genes that are significantly differentially expressed by enhancer silencing.

Principal component analysis showed that samples cluster both by LPS condition and gRNA (Fig. 2b), indicating that both promoter and enhancer silencing have strong effects on the transcriptome. *IRF1* expression in CRISPRi controls shows the expected upregulation with LPS (Wilcoxon p = 0.024 at 90mins and p = 0.038 at 12h compared to 0h), and promoter silencing shows strong repression of *IRF1* (Fig. 2c; Wilcoxon p = 7.4−10^−7^ across time points), indicating a successful repressive effect of the promoter gRNA. The effect of the enhancer silencing is not significantly different from the controls for *IRF1* (Wilcoxon p = 0.8) or for other genes in the locus (P > 0.1) (Supplementary Figure 4). However, given that the principal component analysis indicates a transcriptome-wide effect of the enhancer silencing, we hypothesized that we may be underpowered to detect an effect on *IRF1* in *cis*, but might capture its *trans* effects.

Thus, we performed transcriptome-wide differential expression analysis to detect the transcriptional effect of LPS timepoints, gRNAs and the interaction of these. First, we verified a robust LPS response in the CRISPRi controls, with 30 differentially expressed genes at 90 minutes (FDR 5%) and 476 genes at 12h compared to the 0h samples (Supplementary Table 3), with both gene sets being enriched for GO categories related to inflammatory response (Supplementary Table 4). Next, analyzing the promoter and enhancer silencing effect compared to controls at 12h, we detected 1016 differentially expressed genes for the promoter and 225 genes for the enhancer silencing (Supplementary Table 3), both having an enrichment of many cellular processes including pathways related to immune response, LPS binding and TLR4 activation (Supplementary Table 4). The differential expression of promoter and enhancer have a strong positive correlation (rho = 0.89, p< 2.2e-16, Fig. 2d), which indicates that enhancer silencing downregulates *IRF1*. The enhancer silencing effect was stronger under LPS stimulus than under baseline (Supplementary Figure 5), suggesting that the enhancer is activated under immune response. Neither enhancer nor promoter differentially expressed gene lists were significantly enriched for msigdb IRF1 targets (Fisher test p = 1 and 0.245, respectively), possibly because differentially expressed genes are expected to include direct and indirect targets and downstream pathway effects of IRF1. Altogether, these results show that repression of the enhancer affects the transcriptome, indicating that it is an active regulatory region, and the response being similar to the *IRF1* promoter silencing further provides strong evidence that it is indeed an enhancer of *IRF1*.

### Detection of variant effects on gene expression by genome editing

Enhancer silencing does not necessarily reflect the effect that a genetic variant has on gene expression in *cis* and *trans*. Thus, in order to test whether rs17622517 is indeed the causal eQTL variant and characterize its effect, we used CRISPR/Cas9 genome editing to introduce indels at the variant’s genomic locus in HEK293/TLR4 cells (Fig 3a); SNP editing in the locus with homology-directed repair had too low efficiency to be applicable. From a population of edited cells with 50% non-homologous end joining (NHEJ) based on targeted sequencing, we isolated and grew monoclonal cell lines, and genotyped them by sequencing. We focused on 1-20 bp deletions, allowing a mix of different edited alleles in a given clone, and because the HEK293 cell line is triploid at this locus, clones were considered heterozygous if they have one or two edited alleles. Fifteen clones with initially promising genotypes were further analyzed by long-range PCR to exclude one clone with a large rearrangement at the locus (Kosicki et al., 2018) (Supplementary Figure 6A) and one heterozygous clone was excluded due to a 50/50 allelic ratio suggesting that one allele had been lost. The 13 clones selected for functional follow-up include five homozygous reference (wild type) clones, three heterozygous clones, and five homozygous deletion clones (Supplementary Figure 6B). Each of the 13 clones was exposed to LPS as above with 0h, 90 min, and 12h LPS treatment, after which RNA was extracted and sequenced, and gene expression was quantified. This was done in two replicates in order to reduce any potential noise especially in the stimulus step, and the reads from the replicates were combined for each sample, resulting in a median of 31 million reads per sample.

**Figure 3.**
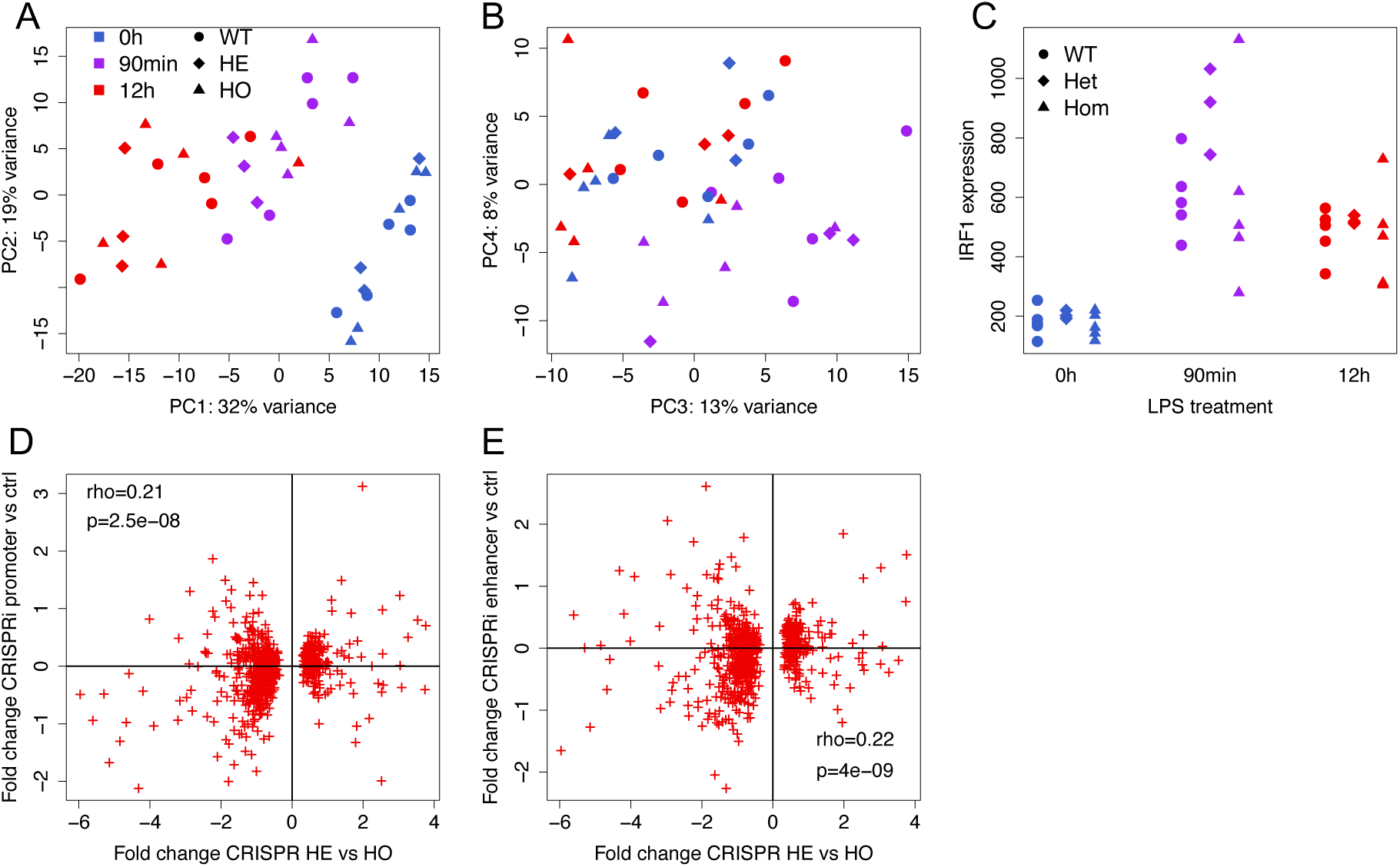
Genome editing with CRISPR. a-b) principal component analysis of gene expression for the different genotypes and LPS conditions, with principal components 1 and 2 in (a) and 3 and 4 in (b); c) IRF1 expression (gene counts normalized to sample read depth) for the different genotypes and conditions; d-e) Correlation of gene expression fold changes at 12h LPS condition for genes that are significantly differentially expressed in heterozygous lines compared to homozygous lines (x-axis), versus expression fold change for promoter (d) and enhancer (e) silencing (y-axis).

The samples cluster strongly by LPS treatment in principal component analysis, as expected, with a potential slight genotype effect (Fig 3a,b). Differential expression analysis (Supplementary Table 5) confirmed the widespread LPS stimulus effect on gene expression of relevant pathways (Supplementary Table 6). *IRF1* expression is induced upon stimulation at 90m and 12h compared to the controls, but it is not significantly differentially expressed between genotypes (log2 fold change 0.37, p = 0.29 heterozygous vs wild-type, log2 fold change −0.03, p = 0.93 homozygous vs wild-type, log2 fold change −0.40, p = 0.26 homozygous vs heterozygous, Fig 3c), and while there are some suggestive patterns especially at 90 mins, they do not appear to recapitulate the *cis*-eQTL pattern. This may be due to low power with a limited number of clones, indels at the locus having a different effect than the rs17622517 SNP, and the cell lines potentially having other on-target or off-target mutations not detected by our genotyping assays. However, again hypothesizing that genetic perturbations at this locus could have downstream effects that we are underpowered to see for *IRF1*, we next analyzed transcriptome-wide differential gene expression, using an interaction model between genotype and condition that would capture edited clones differing in their transcriptional response to LPS stimulation. At 5% FDR we can detect only up to 37 differentially expressed genes (Supplementary Table 5) with no apparent link to IRF1 targets (Supplementary Table 6). Next, we inspected whether the genetic mutations elicit a similar transcriptome effect at 12h as *IRF1* silencing with CRISPRi silencing. The effect of the genetic mutations appear to be best captured by comparison of homozygous and heterozygous clones (762 differentially expressed genes at 25% FDR, < 10 for other genotype comparisons). Using this set of genes, we discovered a significant correlation of the fold changes with those from both promoter (rho=0.21, p=2.52e-8) and enhancer (rho=0.22, p=4.31e-9) silencing (Fig. 3d and e). This suggests that genetic mutations at the rs17622517 locus are causal variants affecting the transcriptome via regulation of *IRF1*, even though we are underpowered to detect effects on individual genes in *cis* and largely also in *trans*.

### Trans-eQTL effects in CRISPRi and CRISPR models

Having provided experimental evidence that rs17622517 is the causal eQTL variant, located in an enhancer that affects transcriptional immune response via *IRF1*, we sought to analyze whether the observed *trans*-eQTL effects in primary monocytes can be recapitulated in this experimental model. Thus, we analyzed trans-eGenes at a relaxed 50% FDR, and correlated their trans-eQTL effect sizes with expression fold change in CRISPRi and CRISPR experiments, all under the late LPS treatment. Correlations for all the comparisons are shown in Fig 4c and Supplementary Figure 7. We observed a significant correlation of *trans*-eQTL effects with differential expression in the CRISPR cell lines for wild type/heterozygote (rho = 0.32, p = 3.7e-5, Fig. 4a) and heterozygote/homozygote (rho = −0.27, p = 5.2e-4, Supplementary Figure 7B) comparisons. The opposing direction of the correlations might be explained by the suggestive pattern of IRF1 expression of the genotype classes (Fig. 3c), with the heterozygous clones having upregulation and homozygotes having downregulation of IRF1, compared to wild type at 90 minutes. Altogether, the significant correlations provide evidence that the trans-eQTL is a true association that can be captured in our cellular model (Fig 4d). We observed no correlation of the trans-eQTL effects with CRISPRi enhancer or promoter silencing (Fig. 4b,c). This suggests that even though the analyses above indicate that both enhancer silencing and genome editing disrupt enhancer function, CRISPRi is suboptimal for recapitulating eQTL effects driven by genetic variants.

**Figure 4.**
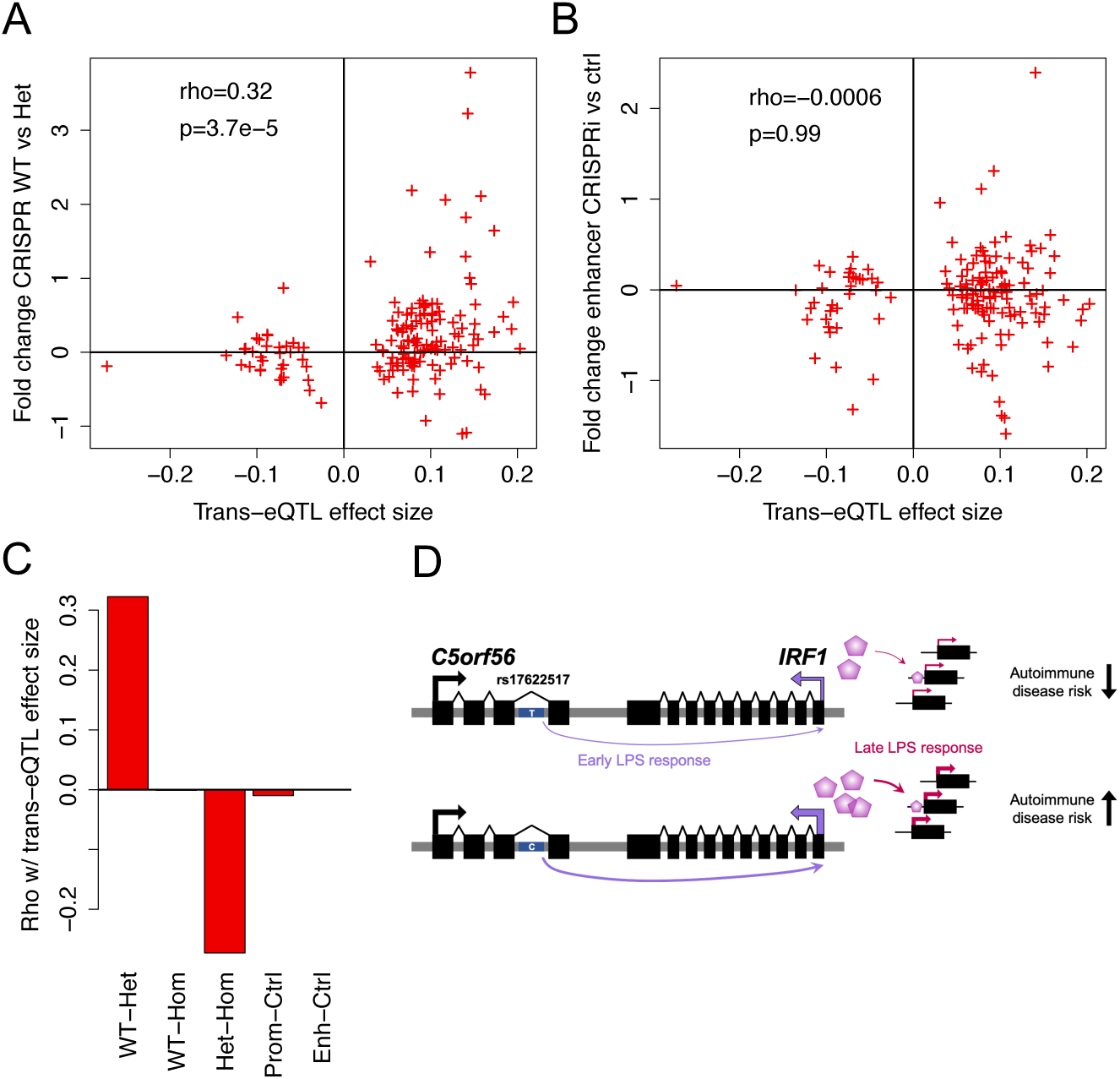
Trans-eQTL effects captured by experimental data. a) Correlation of trans-eQTL effect size with differential expression in CRISPR-edited heterozygous clones, b) after silencing of the enhancer locus with CRISPRi, and c) summarized for all the experimental comparisons. d) Summary of the model supported by data of this study, with the rs17622517 C allele increasing the expression of IRF1 in cis under early immune stimulation, with downstream effects in trans under late immune stimulation. The C allele is also a risk allele for several autoimmune traits.

## Discussion

In this study, we discovered a condition-specific monocyte eQTL that affects the expression of *IRF1* in *cis* and many genes in its pathway in *trans*, so that alternative allele carriers have an increased upregulation of *IRF1* under immune stimulus. This is a plausible molecular mechanism for inflammatory bowel disease associations in this locus, with genetic predisposition for overactivation of *IRF1* immune response (by the rs17622517 C allele) likely contributing to autoimmune disease risk. This further strengthened by two independent eQTLs for *IRF1* having a GWAS signal for autoimmune traits.

We took a CRISPR-based approach to characterize this association in detail in a cellular model. Using a combination of enhancer and promoter silencing with CRISPRi and genome editing, we showed that the causal regulatory variant is rs17622517, affecting the activity of an enhancer located in an intron of *C5ORF56* with a cis-regulatory effect on *IRF1* under immune stimulation with LPS. The effect on the expression of downstream genes in *trans* recapitulates the trans-eQTL association and shows how a genetic effect on a transcription factor such as *IRF1* can affect entire downstream pathways.

Our work highlights both the power and challenges of CRISPR-based experimental analysis of functional effects of genetic variant. The eQTL mapping in primary monocytes under different treatments allowed us to build strong hypotheses of the mechanism that made the experimental follow-up practically feasible, demonstrating how observational functional genetics data helps to guide experimental work. Even though our modified HEK293 cell line is an imperfect model for events observed in primary monocytes, we were able to capture trans-eQTL effects in our cellular system. However, much more data will be needed to build a generalizable understanding of how well functional genetic effects can be extrapolated between cells in their physiological environment and *in vitro*. Our results also highlight the limitations of CRISPR genome editing for functional follow-up: HDR efficiency is too low in many loci, including ours, for feasible analysis of specific SNPs, and interpretation of effects of different indels can be difficult. Perhaps most importantly, the undesired and often undetected genetic or epigenetic differences between the clones introduce a large amount of variation, which makes well-powered and robust analysis difficult without a very large number of clones from each genotype. A key approach in this study to mitigate the limitations of genome editing was to complement it with CRISPRi silencing. Furthermore, while both CRISPR and CRISPRi analyses had limited power to detect robust effects on *IRF1* expression in *cis*, RNA-sequencing allowed us to leverage the power of the entire transcriptome, substantially improving our sensitivity to detect subtle regulatory effects.

The genetic effect on *IRF1* and downstream gene expression in this locus manifests only under immune stimulus, and thus an individual’s transcriptional response to immune stimulus partially depends on the rs17622517 genotype. Such gene-environment interactions are common in *cis* (Barreiro et al., 2012; Fairfax et al., 2014; Kim-Hellmuth et al., 2017; Lee et al., 2014; Nédélec et al., 2016), but few condition-specific master *trans*-regulatory variants such as the one characterized here have been well characterized before (Fairfax et al., 2014; Quach et al., 2016). The association of this locus to inflammatory bowel disease, the known link between the IRF1 pathway and IBD (de Lange et al., 2017; Ellinghaus et al., 2016; Huang et al., 2017), and the link between *IRF1* variants and eczema (Kichaev et al., 2019) and asthma (Demenais et al., 2018; Ferreira et al., 2017) indicate that genetic *IRF1* dysregulation affects disease risk. The specific mechanistic link between the transcriptional IRF1 response described here and autoimmune disease is unknown. However, *IRF1* is known to induce expression of type-I interferons (Miyamoto et al., 1988) and pro-inflammatory cytokines, which can trigger an inflammatory response in a broad range of cell types (Kröger et al., 2002). This is a likely mechanism for *IRF1*’s role in autoimmune disease and a potential topic to be explored in future research. The role of *IRF1* in TLR4 signaling in response to LPS has not been well characterized (Kim-Hellmuth et al., 2017; Pan et al., 2013), and our finding indicates that *IRF1* plays a role in LPS response in the innate immune system. However, the effects described here and their relation to disease mechanism are not necessarily specific to LPS stimulation in monocytes, but could potentially be active in a wide range of immune cells and stimuli.

Altogether, our integrated analysis of genetic effects on gene expression in primary cells and in experimental models provides evidence that within the autoimmune disease risk locus near IRF1, rs17622517 is a causal variant that affects *IRF1* expression in monocytes under LPS stimulation and leads to overactivation of its downstream regulatory pathway.

## Methods

### eQTL discovery

The *IRF1 cis* response eQTLs (reQTLs) were discovered in a previous immune-response eQTL study(Kim-Hellmuth et al., 2017). Briefly, primary monocytes were isolated from 134 donors and treated with LPS, MDP, IVT RNA, or no treatment. To assess early and late immune response, RNA expression was measured with a microarray chip after 90 min and 6 h. To detect a significant *cis*-reQTL, the effect size of association between genotype and expression of genes were compared between untreated and treated samples. rs17622517 was found to be a significant *cis-*reQTL variant for *IRF1* at 90m after LPS treatment, i.e. under early immune stimulus.

*Trans-*eQTL discovery was done with the MatrixEQTL R package, performing a linear regression between the genotype of the top variant for each of the 126 significant *cis-*reQTL at LPS 90 min and all expressed probes in monocytes at 6 hours (22,657). Benjamini-Hochberg correction was performed across p-values for all variants and probes. A significance threshold of FDR < 0.5 was used for downstream analyses. Since rs17622517 was found to be associated with many genes in *trans* at 6 hours, in addition to the *cis* association with *IRF1* and GWAS association for autoimmune diseases, it was selected for further follow up.

Effect size of trans-eQTL associations was measured as allelic fold change calculated from linear regression summary statistics (Mohammadi et al., 2017). When multiple expression probes measure the same gene, their mean effect size was used.

### Enrichment of IRF1 targets

*IRF1* targets were obtained from the molecular signatures database (MSigDB) gene set *IRF1*-01, which comprises genes which contain at least one *IRF1* motif in the 4kb upstream and downstream from their transcription start site (Liberzon et al., 2011). *Trans-*eGenes from the rs17622517 *trans-*eQTL (FDR < 0.5) and CRISPRi differentially expressed gene set (FDR < 0.05) were tested for enrichment in *IRF1* target genes using a Fisher’s exact test comparing the number of *IRF1* target genes in these sets compared to the background of all expressed genes.

### Cell culture

HEK293/hTLR4A-MD2-CD14 Cells (Invivogen) were selected for functional follow up of the *IRF1 trans-*eQTL because of the ease of transfection of HEK293 cells and the addition of the TLR4 receptor, which is essential for cellular response to LPS. Cells were cultured in DMEM supplemented with 4.5 g/l glucose (Corning), 10% fetal bovine serum (Sigma-Aldrich), 1% penicillin/ streptomycin (Corning) and 1% L-glutaMAX (gibco). Cells were passaged using cell scraping to avoid damaging the cell surface receptors.

### LPS concentration and time point optimization

To determine the optimal LPS concentration and harvesting timepoints for the HEK293-TLR4 cells, we tested their transcriptomic response to LPS under multiple conditions. HEK293-TLR4 cells were plated into three wells in 24-well plates with 180,000 cells per well. 24 hours later, 0 ng, 500 ng, 1000 ng or 2000 ng of LPS (Invivogen) per mL was added to the cells. RNA was extracted at three timepoints for the 500 and 2000 ng conditions (90m, 6h, 24h) and six timepoints for the 1000 ng condition (45m, 90m, 3h, 6h, 12h, 24h), by adding 500 uL of IBI Isolate DNA/RNA Reagent (IBI Scientific) directly to cells on the plate and stored at −80C until extraction.

### RNA extraction and RNA-seq library preparation

RNA was extracted following the Direct-zol RNA MicroPrep kit (Zymo Research) manufacturer’s instructions. RNA was treated with DNAse I (Ambion) and enriched for mRNA using Dynabeads mRNA DIRECT Purification Kit (Thermo Fisher). cDNA was generated using a custom scaled-down modification of the SMART-seq protocol (Picelli et al., 2014). cDNA was synthesized from RNA input using Maxima H Minus Reverse Transcriptase (Thermo Fisher Scientific). It was then amplified using Kapa HiFi 2X Ready Mix (Kapa Biosystems) and cleaned using 0.9X Ampure beads (Beckman Coulter). Finally, cleaned cDNA samples were tagmented and indexed using the Nextera XT DNA Library Prep Kit (Illumina). Library size and tagmentation were confirmed using the TapeStation HS D1000 kit (Agilent). Libraries were pooled in equal molarity and sequenced with the NextSeq 550 High-Output kit (Illumina) with paired-end 75 bp reads.

### RNA-seq data processing

Reads were first trimmed of adapters using trimmomatic, then aligned to the hg19 genome using STAR 2-pass mapping. Gene counts were calculated with FeatureCounts using Gencode v19 gene annotations.

### CRISPRi of IRF1 promoter and enhancer locus

In order to determine whether the region of the eQTL variant regulates IRF1, and the effect of IRF1 perturbation in our cell line, we performed CRISPRi experiments targeting both the variant locus and the IRF1 promoter. HEK293-TLR4 cells were plated in three 24-well plates with 120,000 cells/well in 1 ml of DMEM. 24 hours later, the medium was replaced with 0.5 ml of OptiMem (Gibco) and cells were transfected with 1.5 ul of lipofectamine 3000 (Thermo Fisher Scientific), 200 ng of CRISPR-KRAB-MeCP2 vector (Addgene 110821) and 50 ng of gRNA gblock (IDT) (4 wells each received a gRNA targeting EGFP as a neutral control (‘GGTGGTGCAGATGAACTTCA’), 14 bp downstream of the IRF1 promoter (‘GTCTTGCCTCGACTAAGGAG’) or the enhancer (‘TTCTCTGTAGCCCTTGTATT’)). After 28 hours, 1 ug/mL of LPS (Invivogen) was added to the 90 m and 12 h samples and nothing to the control samples, with four biological replicates of each gRNA for each LPS treatment (36 total samples). Cells were collected at their respective time point with TRIzol reagent (Thermo Fisher Scientific) and stored at −80C before RNA extraction and RNA-sequencing.

### Genome editing

In order to validate the *cis* and *trans* reQTL associations of rs17622517, we edited the HEK293-TLR4 cells using CRISPR/Cas9 genome editing. A gRNA (‘TTCTCTGTAGCCCTTGTATT’) was designed with an NGG PAM and a cut site 1 bp downstream from rs17622517. The gRNA was ordered as a single stranded oligo gblock from IDT and amplified using 2 50 uL reactions of Q5 High Fidelity 2X Master Mix (NEB). Cells were transfected with 0.5 ug gRNA gblock and 2.5 ug px458 plasmid (Addgene plasmid # 48138) containing spCas9 and GFP, using lipofectamine 3000 (Thermo Fisher Scientific). 24-hours later, cells then underwent fluorescence-activated cell sorting for GFP+ cells using a Sony SH800Z cell sorter to enrich for transfected cells. Efficiency of editing was tested using a T7E1 assay and electrophoresis gel to detect presence of NHEJ. The GFP+ cells were also sorted as single cells into 15 96-well plates and expanded into monoclonal cell lines.

Clones were genotyped by creating an amplicon library for each clone from gDNA using nextera primers capturing a 218 bp amplicon containing the variant locus. Indexing PCR was performed using primers specific to the constant sequence on the Nextera primers, resulting in dual-barcoded amplicons with Illumina adapters. Libraries were mixed in equal volume and sequenced on the MiSeq using 150 bp paired-end reads. Fastq files generated by the Illumina software were trimmed for adapter sequences and quality using trimmomatic. Reads were aligned to the genomic locus and categorized as no edit or NHEJ (with indels) using Edityper (Yahi et al. in prep). Since the cell line is triploid at this genomic locus, clones were considered heterozygous if they had NHEJ rate between 20-70%. Clones with less than 10% NHEJ were considered wild type and clones with greater than 90% NHEJ were considered homozygous edited. Five each of wild type, heterozygous and homozygous edited were selected for follow up. BAM files from the selected clones were visually inspected to confirm genotype. In addition, a 3,496 bp PCR followed by electrophoresis gel was performed on the clones in order to check for larger indels not captured by the shorter amplicon libraries.

### LPS treatment of edited clones

To detect the reQTL effect of the edited locus, we performed LPS treatment followed by RNA-sequencing on each isolated edited and wild type cell line. Each clonal cell line was plated into three wells (one well each for control, 90m and 12h treatments) in 24-well plates with 180,000 cells per well. 24 hours later, 1ug/mL of LPS (Invivogen) was added to the 90 m and 12 h samples. RNA was extracted at the designated time point by adding 500 uL of IBI Isolate DNA/RNA Reagent (IBI Scientific) directly to cells on the plate and stored at −80C until extraction and RNA-sequencing. This procedure was done twice to reduce potential technical variability from the LPS treatment and sequencing, and after verifying that the results were generally consistent, the gene count matrices from the two runs were summed together for analysis.

### Differential expression analysis

Differential expression analyses for the CRISPRi and CRISPR data were performed using the R package DEseq2 (Love et al., 2014). We first transformed and normalized the gene count matrices with vst(), and did principal component analysis. Differential expression was performed using interaction models. For CRISPRi we used ∼gRNA + condition + gRNA:condition where gRNA denotes promoter, enhancer or control gRNA, and condition denotes 0, 90 min or 12 h LPS treatment. Additionally, we looked at the promoter vs control and enhancer vs control gRNA effect within 90m LPS samples and 12h LPS samples, and the effects of the LPS treatments within the control samples. For CRISPR, we used a nested interaction model (expression ∼ genotype + genotype:clone + genotype:condition) accounting for the same clone undergoing different LPS treatments. Additionally, we looked at the effects of the two LPS treatments within the WT clones. Prior to p-value correction, genes were discarded if they were not annotated as protein coding or lncRNA or they did not have an average expression of greater than 5 read counts across samples. P-values for differential expression were corrected using Benjamini-Hochberg correction, using FDR< 0.05 threshold.

Enrichment analysis of significantly differentially expressed genes was done using DAVID biological process gene ontology enrichment (Huang et al., 2009), using Benjamini-Hochberg corrected p-values and FDR< 0.05 except for comparison between edited clone and CRISPRi differential expression, where FDR < 0.25 was used.

## Supporting information

Suppementary figures and table legends

## Acknowledgements

This work was supported by National Institutes of Health grants R01MH106842 (T.L.), Roy and Diana Vagelos Precision Medicine Initiative Pilot Grant (T.L.), T32GM008224 (Y.R.), and Marie-Skłodowska Curie fellowship H2020 Grant 706636 (S.K-H.).

## Data availability

The RNA-sequencing fastq files and gene expression matrices are available through NCBI’s gene expression omnibus under series accession number GSE145487.

## Supplementary Materials

Supplementary figures and supplementary table legends are in the enclosed pdf. Supplementary tables are in the enclosed excel file.

