## Supplementary material for "An autoimmune disease risk variant has a *trans* master regulatory effect mediated by IRF1 under immune stimulation": Suppementary figures and table legends

**Supplementary tables description**

**Filename: Supplementary_tables.xls**

**Supplementary Table 1: LPS 6h Trans-eQTLs.** Significant LPS 90 min cis-reQTLs in monocytes were tested against all expressed probes and filtered for an FDR cutoff of 0.5.

**Supplementary Table 2: IRF1 90m cis-eQTL.** Association between genotype of variants within 100 kb of IRF1 transcription start site and IRF1 expression in monocytes treated with LPS for 90 min.

**Supplementary Table 3: Differential gene expression (FDR 0.05) for CRISPRi samples.** hTLR4 Cells were transfected with control, enhancer and promoter gRNAs and a CRISPRi construct. They were then treated with no LPS, LPS for 90 min and LPS for 12h. Differential expression analysis was performed for LPS treatment in CRISPRi control samples, gRNA condition in different LPS treatments, and the interaction between LPS treatment and gRNA condition.

**Supplementary Table 4: Gene ontology enrichment for CRISPRi differentially expressed genes**. Gene ontology enrichment analysis was performed on gene sets from CRISPRi differential expression analysis.

**Supplementary Table 5: Differential gene expression (FDR 0.05) for CRISPR edited clones.** hTLR4 Cells were transfected a CRISPR/Cas9 construct and a gRNA specific for the rs17622517 locus. Wild type, heterozygous or homozygous clones were isolated and then treated with no LPS, LPS for 90 min and LPS for 12h. Differential expression analysis was performed for LPS treatment in WT clones, and the interaction between genotype and LPS treatment.

**Supplementary Table 6: Gene ontology enrichment for CRISPR edited differentially expressed genes**. Gene ontology enrichment analysis was performed on gene sets from CRISPR edited differential expression analysis.

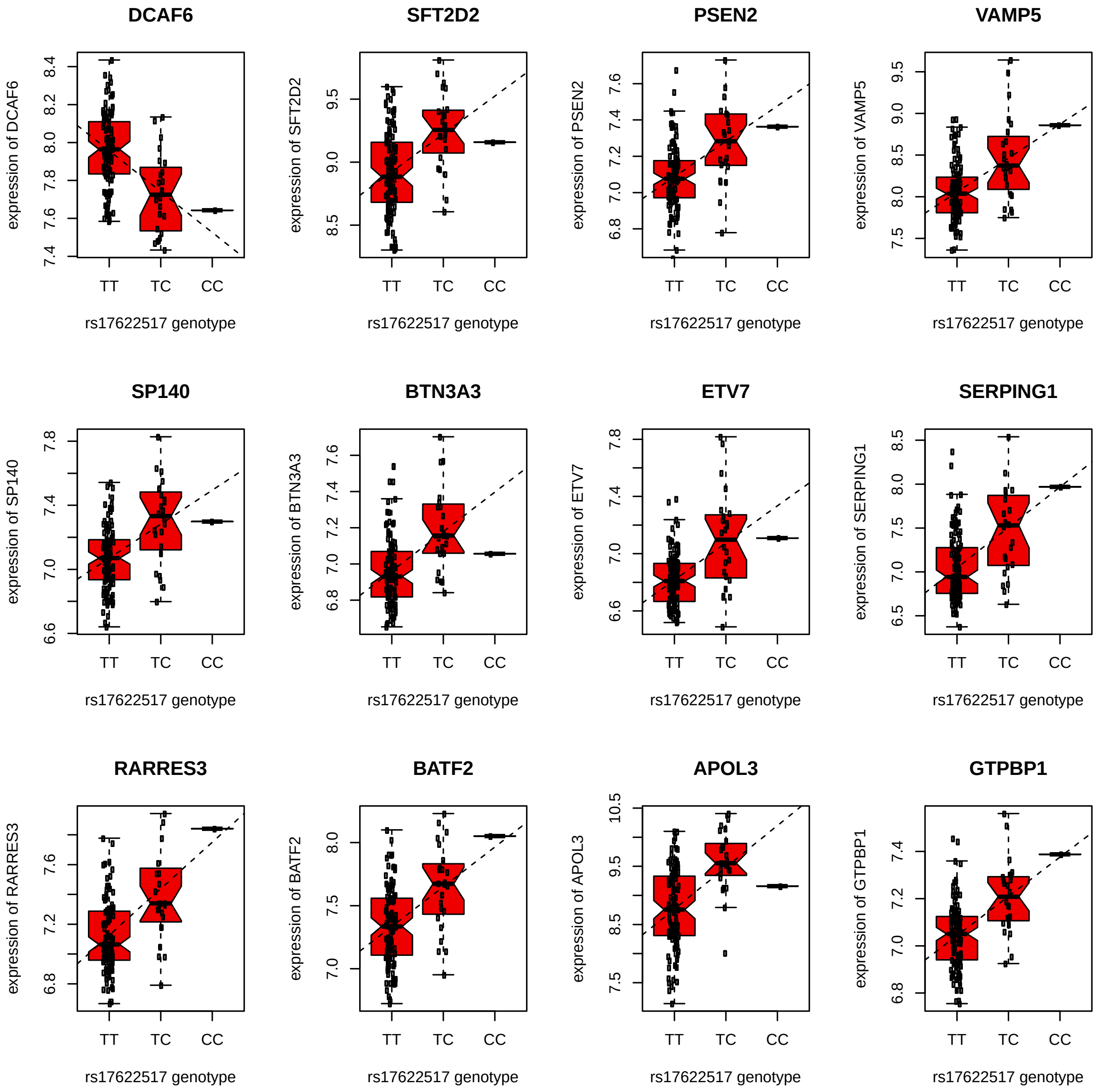


**Supplementary Figure 1. Trans-eQTL associations.** Twelve strongest trans-eQTLs associated with rs17622517 under 6h LPS.

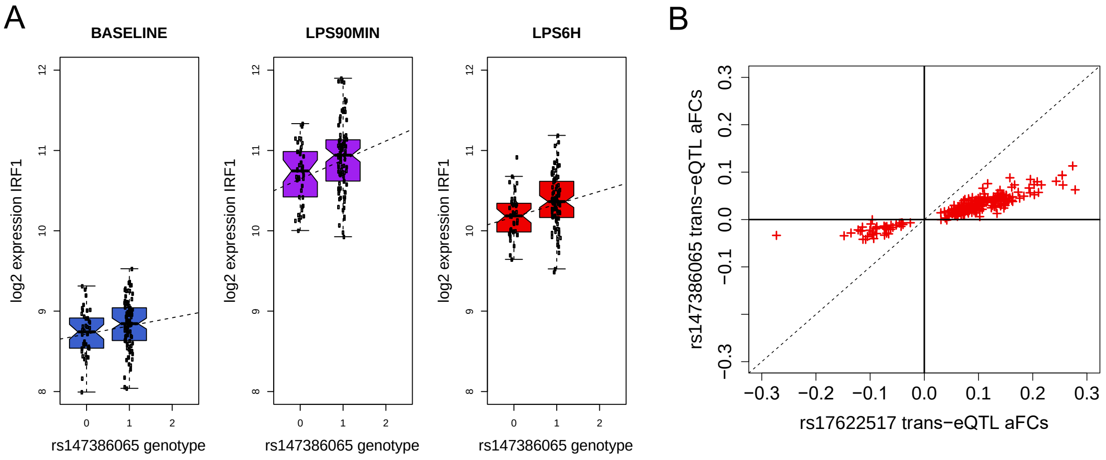


**Supplementary Figure 2. Secondary eQTL signal in the IRF1 locus.** a) cis-eQTL effect of rs147386065 on *IRF1*, and b) *trans*-eQTL effect sizes of the significant rs17622517 trans-eQTLs (FDR < 50%) versus effect sizes of rs147386065 *trans* effects on the same genes.

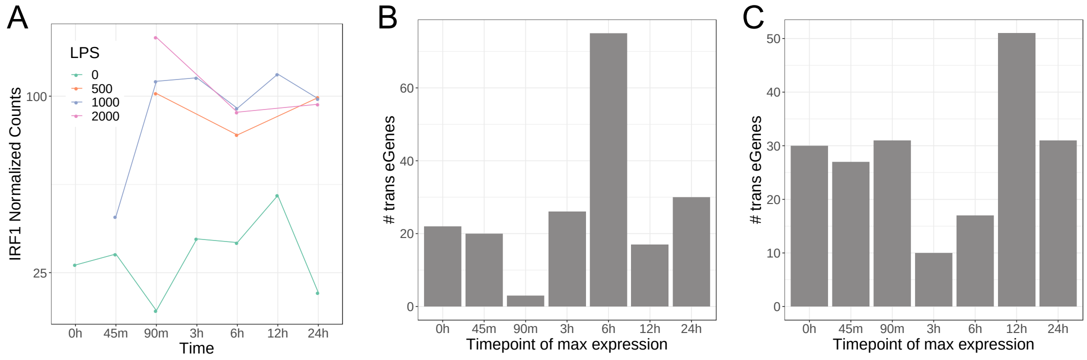


**Supplementary Figure 3. Optimization of the LPS treatment for HEK293-TLR4 cells.** a) expression of *IRF1* for the different conditions*;* b-c) number of rs17622517 trans-eGenes that have their maximum expression at each time point in primary monocytes, using data from Kim-Hellmuth et al. 2017(b), and in HEK293-TLR4 cells (c). The peak trans-eGene expression is at 6 hours in monocytes and at 12 hours in HEK293-TLR4 cells.

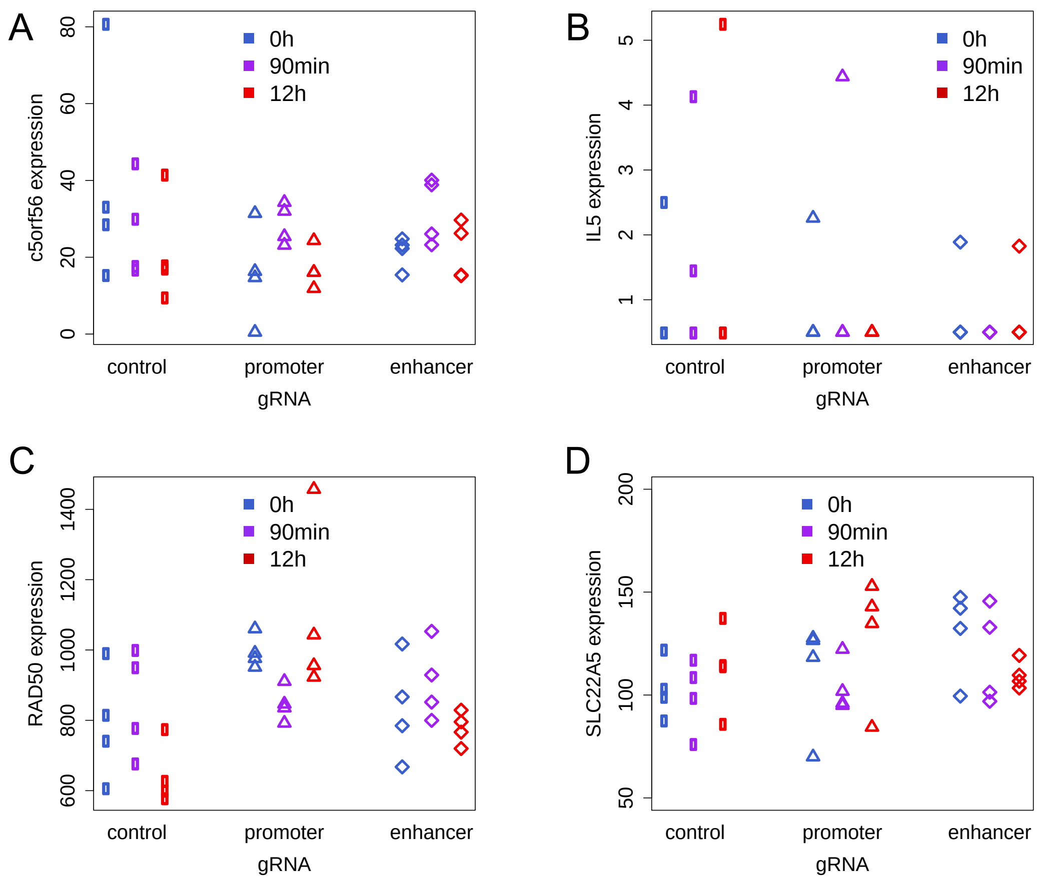


**Supplementary Figure 4. Enhancer silencing effects on gene expression in *cis*.** Expression levels of the genes within 1Mb in HEK293-TLR4 cells with and without CRISPRi silencing of the rs17622517 locus. The lack of significant difference indicates that this putative enhancer does not have a strong effect on other genes in *cis*.

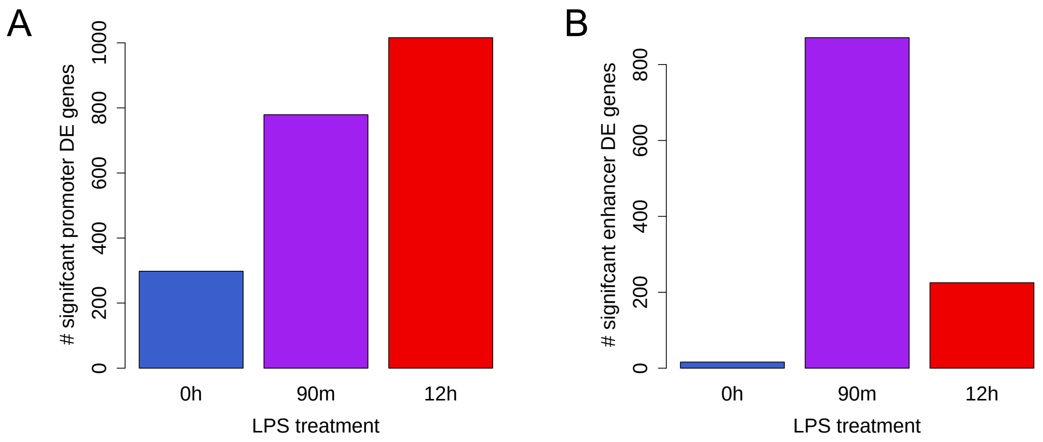


**Supplementary Figure 5. Enhancer and promoter silencing effects for different LPS time points.** Number of significantly differentially expressed genes in promoter (A) and enhancer (b) under different LPS stimulation conditions.

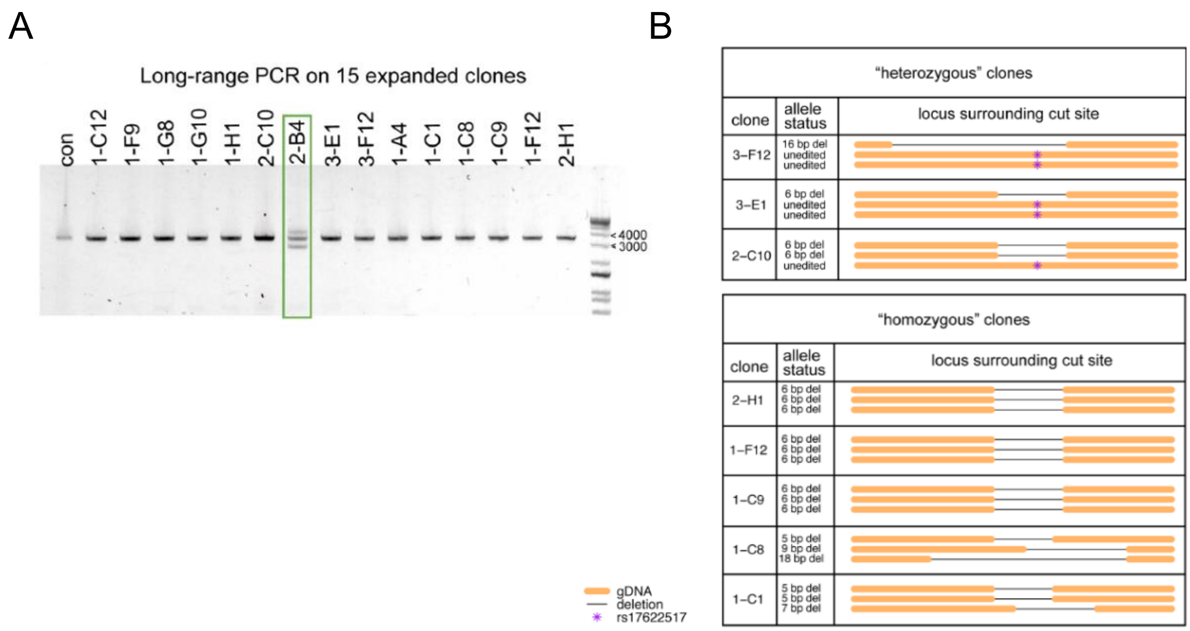


**Supplementary Figure 6. Genotyping of edited clones.** a) Electrophoresis of long-range PCR products showed a large deletion in one clone (green) with that was excluded from further analysis; b) genotypes of clones selected for analysis.

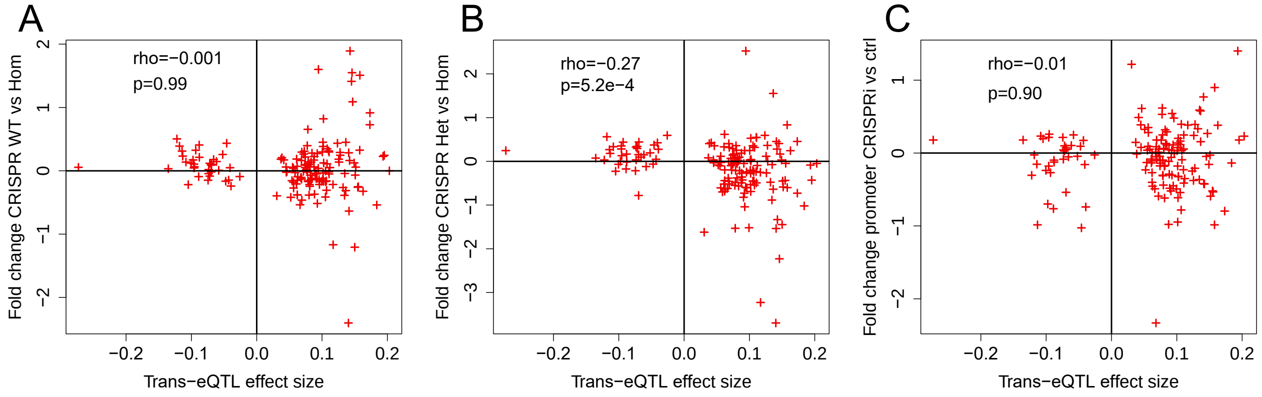


**Supplementary Figure 7. Correlation of trans-eQTL effect sizes and experimental perturbations.** Correlation of trans-eQTL effect size (aFC) with differential expression in a) wild-type versus CRISPR-edited homozygous clones, b) CRISPR-edited heterozygous versus homozygous clones, and c) CRISPRi promoter silencing versus control samples.
